# Levetiracetam treatment normalizes levels of the presynaptic endocytosis machinery and restores non-amyloidogenic APP processing in *App* knock-in mice

**DOI:** 10.1101/2021.02.22.432282

**Authors:** Nalini R. Rao, Jeffrey N. Savas

## Abstract

Increasing evidence indicates that toxic amyloid-beta (Aβ) peptides, produced by sequential proteolytic cleavage of the Amyloid Precursor Protein (APP), induce neuronal circuit hyperexcitability in the early stages of Alzheimer’s disease (AD). As a result, treatments that modulate this early excitatory/inhibitory imbalance could act as potential AD therapies. Levetiracetam, an atypical antiepileptic drug, has garnered recent interest, despite the mechanism(s) of action remaining elusive. In this study, we set out to identify the pathways and mechanisms primarily affected by levetiracetam in diseased brains of amyloid pathology. Using the *App* knock-in mouse models and multiplexed TMT-quantitative mass spectrometry-based proteomic analysis to determine how levetiracetam affects the proteome, our findings demonstrate that levetiracetam treatment selectively normalizes levels of presynaptic endocytosis proteins and is capable of lowering Aβ_42_ levels by altering APP processing. These novel findings demonstrate a mechanism of action for how levetiracetam lowers Aβ_42_ production.

## INTRODUCTION

In Alzheimer’s disease (AD), sequential proteolytic cleavage of the Amyloid Precursor Protein (APP) leads to the production of toxic amyloid-beta (Aβ) peptides. The inability to efficiently degrade Aβ_42_ has been shown to drive downstream pathologies such as synapse deterioration and the formation of amyloid plaques and neurofibrillary tangles (Zhou et al., 2015, Lacor et al., 2007, Lambert et al., 2006, Herrup 2015). While downstream repercussions have been documented, there is currently no effective treatment to prevent, reverse, or slow the progression of AD. This is most likely due to the Aβ lowering antibody treatments being administered too late in the progression of AD (Herrup 2015). As a result, there has been a shift in focus to investigating and identifying early AD pathologies that may serve as potential therapeutic targets.

Hyperactivity and neural network disruptions have been observed during the initial stages of amyloid pathology and could represent a pioneering aspect of AD pathogenesis (Palop et al., 2009, Kuchibhotla et al., 2008, Palop et al., 2007). These findings have motivated recent investigations focused on the role of brain hyperexcitability in AD and subsequently whether modulating the excitatory/inhibitory imbalance could be an efficacious AD therapy. We recently discovered an early impairment in degradation and turnover of presynaptic synaptic vesicle (SV) machinery proteins in the APP knock-in (*App* KI) mouse models of amyloid pathology that precedes Aβ_42_ accumulation (Hark et al., 2020). Our findings indicate that targeting or correcting early presynaptic dysfunction could represent an effective therapeutic target. Levetiracetam is an atypical antiepileptic drug that, unlike those targeting the GABA-ergic system, binds to the presynaptic synaptic vesicle glycoprotein 2A (SV2A) (Vossel et al., 2017). However, despite FDA approval and wide use, levetiracetam’s mechanism(s) of action remain elusive. In mouse models of AD-like pathology, levetiracetam administration reduces hyperexcitability, suppresses neuronal network dysfunction, and decreases Aβ plaque burden and the corresponding cognitive deficits (Vossel et al., 2017, Sanchez et al., 2012, Shi et al., 2013, Koh et al., 2010, Nygaard et al., 2015). Levetiracetam is, to date, the focus of seven Phase 1 or 2 clinical trials for AD (Toniolo et al., 2020).

In this study, we set out to identify the pathways and mechanisms primarily affected by levetiracetam in diseased brains of amyloid pathology to determine how levetiracetam affects the proteome. In order to avert the possible confounding effects of APP overexpression, we used *App* KI mouse models that have the humanized Aβ_42_ sequence expressed under the endogenous APP promoter and harbor familial mutations (Saito et al., 2014). Here, we show that chronic levetiracetam administration decreases cortical Aβ_42_ levels and lowers the amyloid plaque burden. In addition, using multiplexed TMT-quantitative mass spectrometry-based proteomic analysis, we determined that chronic levetiracetam administration selectively normalizes levels of presynaptic endocytic proteins. Finally, we found that levetiracetam treatment selectively lowers β-CTF levels while the abundance of full-length APP remains unchanged. In summary, this work reports that chronic treatment with levetiracetam serves as a useful therapeutic in AD by normalizing levels of the presynaptic endocytic proteins and altering APP cleavage preference leading to a decrease in both Aβ_42_ levels and the amyloid plaque burden. These novel findings provide pioneering evidence for the previously documented therapeutic value of levetiracetam in mitigating AD pathology.

## RESULTS

### Synaptic vesicle-associated proteins have altered abundance in *App*^*NL-F/NL-*F^and *App*^*NL-G-F/NL-G-F*^ cortical extracts

We deigned our experiments to investigate the effect of chronic levetiracetam in *App* KI brains with varying degrees of Aβ_42_ pathology. The *App*^*NL/NL*^ model serves as a relative control that does not develop Aβ_42_ pathology, while *App*^*NL-F/NL-F*^ mice have relatively slow progressing Aβ_42_ pathology. *App*^*NL-G-F/NL-G-F*^ present with aggressive Aβ_42_ pathology that is abundantly present by six months of age (Fig. 1a). Levetiracetam at 75 mg/kg or vehicle saline solution were administered intraperitoneally daily for 30 days beginning at six months for each *App* KI model (Fig. 1a). To investigate levetiracetam’s mode of action, we performed a quantitative bottom-up proteomic screen using cortical extracts with 16-plex tandem mass tag (TMT) mass spectrometry (TMT-MS). We compared protein abundance between cohorts given vehicle (N = 6) or levetiracetam (N = 6) of the same *App* KI genotype along with multiple float channels that allow for comparisons between the multiple TMT-MS experiments (i.e. genotypes). The overall TMT channel peak intensities were similar in all three experiments, indicating efficient labeling (Fig. 1b). To assess the reliability of the TMT-MS data, we plotted the number of total quantified proteins, reporter ion intensities, and fold change distribution, and confirmed similar data quality (Fig. S1a-d). We compared the protein abundance in *App*^*NL-F/NL-F*^ to *App*^*NL/NL*^ and identified 1,619 significantly altered proteins (Fig. 2a-c; Table S1). In the parallel *App*^*NL-G-F/NL-G-F*^ dataset, we identified 1,266 significantly altered proteins compared to *App*^*NL/NL*^ (Fig. 2a-c; Table S1). To mine the significantly regulated proteins in *App*^*NL-F/NL-F*^/*App*^*NL/NL*^ and *App*^*NL-G-F/NL-G-F*^/*App*^*NL/NL*^, we performed Gene Ontology (GO) cell component (GO:CC) enrichment analysis with PANTHER. In both datasets, the regulated proteins are significantly enriched for the GO:CC terms: synaptic vesicle (SV), synapse, presynapse, postsynapse, and others (Fig. 2d-e; Table S2). This is consistent with our previous report that *App*^*NL-F/NL-F*^ and *App*^*NL-G-F/NL-G-F*^ brains both have synaptic proteome alterations by six months of age [Hark et al., 2020]. To further explore these two datasets, we extracted the abundance of a panel of SV associated proteins that we previously found elevated due to impaired degradation and found highly similar trends (Hark et al., 2020). (Fig. 2f). These results confirm the reliability of our TMT-MS analyses and extend our previous findings that the axon terminal proteome represents an early site of amyloid pathology.

**Figure 1.**
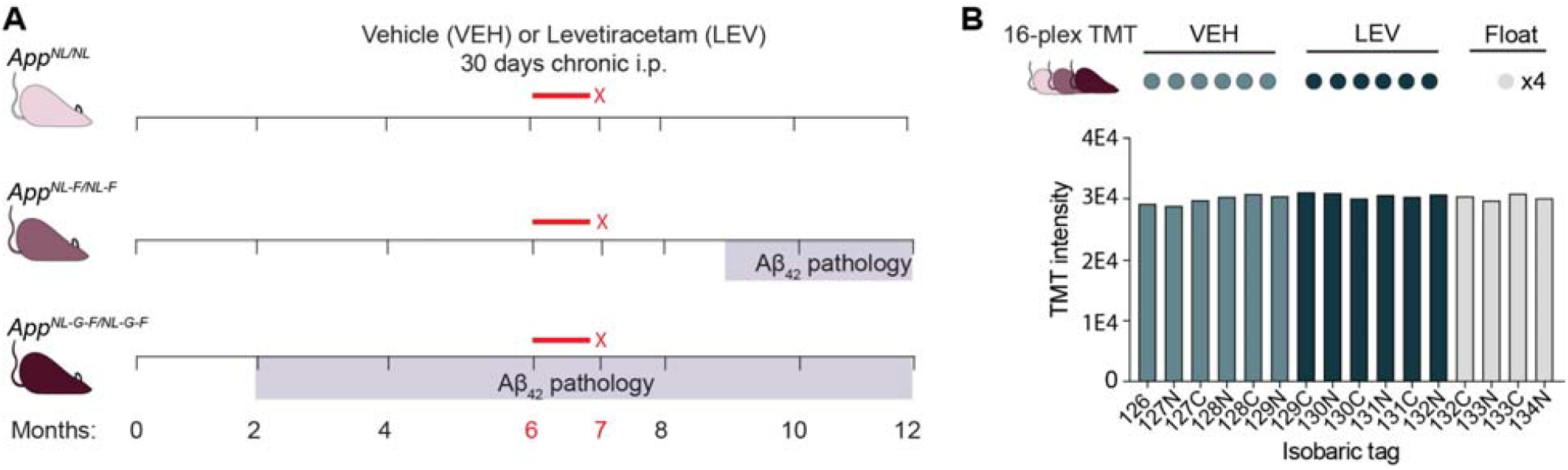
Timeline of amyloid pathology and chronic levetiracetam treatment in *App* KI mouse models. (**A**) Schematic depicting drug injections in relation to the onset of Aβ_42_ pathology in *App*^*NL/NL*^, *App*^*NL-F/NL-F* F^, and *App*^*NL-G-F/NL-G-F*^ genotypes. Mice from each of the three *App* KI genotypes were vehicle (VEH) (N=6) or levetiracetam (LEV) treated (N=6). (**B**) Schematic depicting the 16-plex TMT-MS experimental design. Each genotype, *App*^*NL/NL*^, *App*^*NL-F/NL-F*^, and *App*^*NL-G-F/NL-G-F*^ was analyzed in a 16-plex TMT-MS experiment comparing VEH to LEV treatments with four float channels for data normalization between experiments. Representative overall TMT channel peak intensities for each isobaric tag from the *App*^*NL-G-F/NL-G-F*^ 16-plex TMT-MS experiment demonstrate equal labelling across all channels.

**Figure 2.**
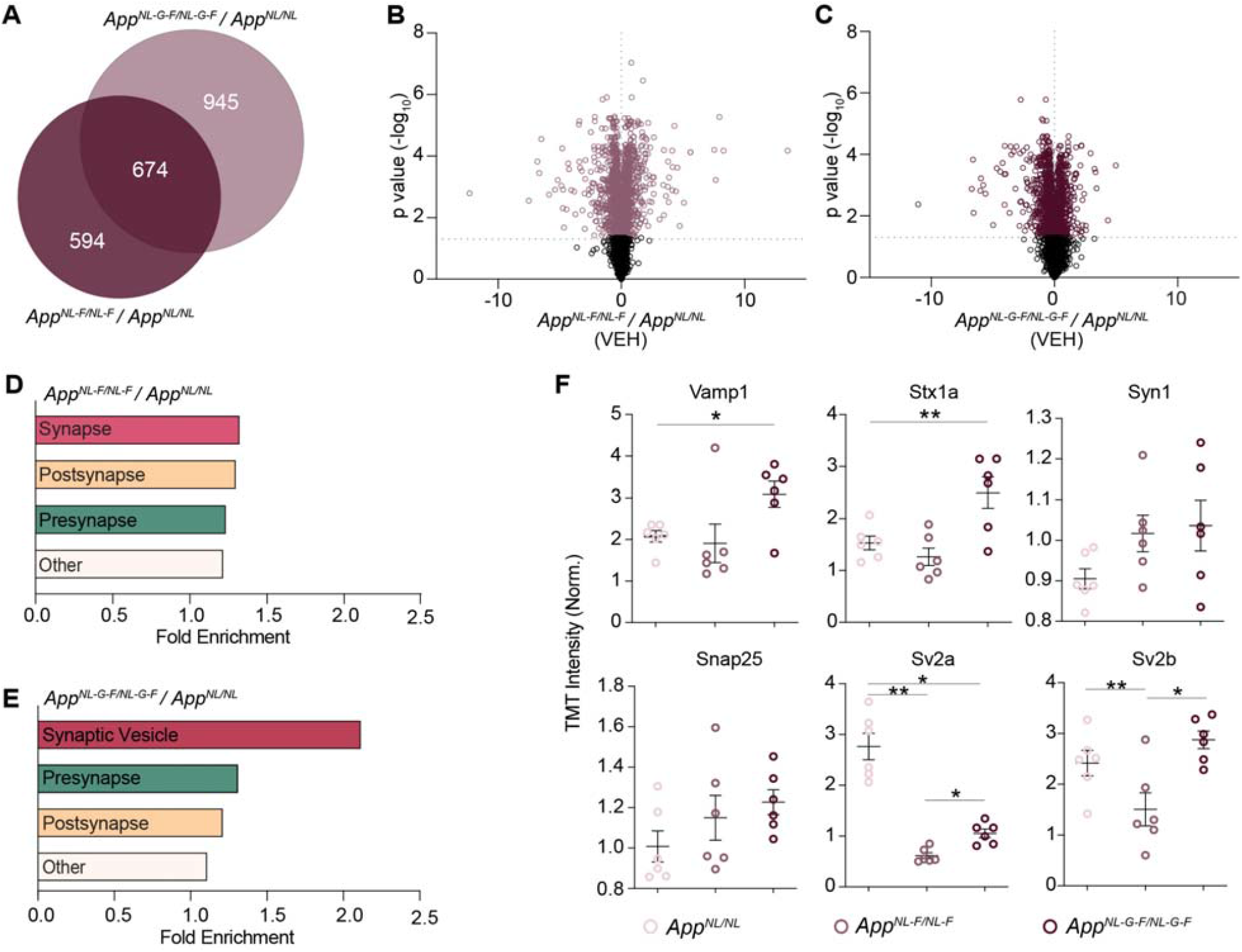
Synaptic vesicle machinery proteins have selective and significantly altered fold change in *App*^*NL-F/NL-F*^/*App*^*NL/NL*^ and *App*^*NL-G-F/NL-G-F*^/*App*^*NL/NL*^ cortical extracts. (**A**) Venn diagram of significantly altered proteins (B.H. p-value < 0.05) between *App*^*NL-F/NL-F*^ /*App*^*NL/NL*^ and *App*^*NL-G-F/NL-G-F*^ /*App*^*NL/NL*^. Values indicate total number of significantly altered proteins. (**B, C**) Volcano plots depicting protein fold change for vehicle *App*^*NL-F/NL-F*^ compared to *App*^*NL/NL*^ (B) and vehicle *App*^*NL-G-F/NL-G-F*^ compared to *App*^*NL/NL*^ (C). Significant proteins (B.H. p-value < 0.05) are colored and non-significant proteins are shown in grey (Table S1). (**D, E**) Gene ontology (GO) cell component enrichment (GO:CC) analysis of significantly altered proteins in vehicle *App*^*NL-F/NL-F*^ / *App*^*NL/NL*^ (D) and vehicle *App*^*NL-G-F/NL-G-F*^ / *App*^*NL/NL*^ (E) (Table S2). Bar graphs depict fold enrichment of each significant GO term. (**F**) Representative normalized TMT intensities across genotypes for a panel of selected synaptic vesicle proteins. N=6 for each group. Data represents mean ± SEM, analyzed with unpaired Student’s t-test or one-way ANOVA with post-hoc Sidak test. * = p-value <0.05, ** = p-value < 0.01.

### Chronic levetiracetam administration normalizes levels of presynaptic endocytic proteins in *App*^*NL-G-F/NL-G-*F^cortex

To investigate the effect of chronic levetiracetam on Aβ_42_ levels, we extracted the relative APP peptide abundance mapping either inside or outside the Aβ_42_ amino acid sequence. Notably, in cortical extracts from levetiracetam treated *App*^*NL-G-F/NL-G-F*^ mice, peptides mapping to the Aβ_42_ amino acid sequence were significantly reduced compared to vehicle treated controls (p-value = 0.0255) (Fig. 3a). As expected, in levetiracetam versus vehicle treated *App*^*NL/NL*^ mice, Aβ_42_ peptides showed no significant difference (p-value = 0.4960) (Fig. 3a). On the other hand, APP peptides mapping outside of Aβ_42_ showed no difference in abundance between levetiracetam versus vehicle in both *App*^*NL/NL*^ and *App*^*NL-G-F/NL-G-F* F^ [F(3,20) = 2.717, p-value = 0.0719] (Fig. 3b). Overall we found that chronic levetiracetam had no significant effect on global protein abundance [F(3,20) = 1.988, p-value = 0.1483] (Fig. 3c). These findings indicates that levetiracetam treatment can lower steady state Aβ_42_ levels without altering overall APP levels in *App*^*NL-G-F/NL-G-F*^ mice, which at six months of age harbor extensive Aβ_42_ pathology.

**Figure 3.**
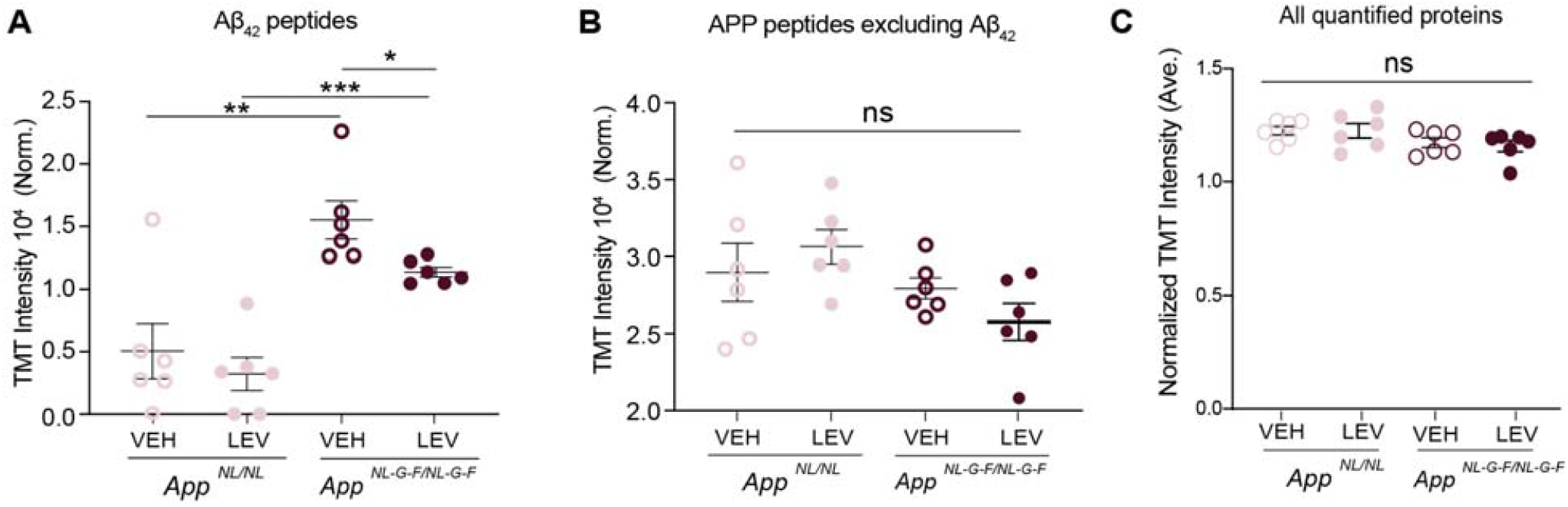
Chronic levetiracetam administration selectively lowers levels of Aβ_42_ in *App*^*NL-G-F/NL-G-F*^ cortex. (A) Normalized TMT intensities of APP peptides within the Aβ_42_ sequence comparing VEH and LEV groups of *App*^*NL-G-F/NL-G-F*^ and *App*^*NL/NL*^ animals. (**B**) Normalized TMT intensities of APP peptides excluding those within the Aβ_42_ amino acid sequence comparing VEH and LEV groups of *App*^*NL-G-F/NL-G-F*^ and *App*^*NL/NL*^ animals (**C**) Bar graph depicts normalized global TMT intensities for all proteins in *App*^*NL-G-F/NL-G-F*^ and *App*^*NL/NL*^ VEH and LEV groups.

We next sought to investigate how levetiracetam affects the *App*^*NL-G-F/NL-G-F*^ cortical proteome. We homed in on the proteins that were significantly altered between the vehicle treated cohorts of *App*^*NL-G-F/NL-G-F*^ and *App*^*NL/NL*^. Next, we probed proteins from this comparison that were normalized by levetiracetam treatment (i.e. genotype vs drug effect). Of the 1,578 significantly altered proteins in vehicle *App*^*NL-G-F/NL-G-F*^/*App*^*NL/NL*^, 985 of those proteins were no longer significantly altered in the levetiracetam *App*^*NL-G-F/NL-G-F*^/*App*^*NL/NL*^, indicating that their levels were modulated by levetiracetam (Fig. 4a; Table S3). We then performed GO:CC enrichment analysis of the 985 proteins with PANTHER, and found that normalized proteins are most significantly enriched for GO:CC terms: presynaptic endocytosis, postsynapse, and synapse, among others (Fig. 4b; Table S4). We focused on the proteins belonging to the GO:CC term presynaptic endocytosis and investigated how their levels changed with treatment by comparing vehicle to levetiracetam datasets for *App*^*NL-G-F/NL-G-F*^. Notably, nearly all of the presynaptic endocytosis proteins had elevated levels after levetiracetam treatment (Fig. 4c). We found that levetiracetam treatment in *App*^*NL-G-F/NL-G-F*^ normalized presynaptic endocytosis protein levels back towards *App*^*NL/NL*^ control levels. (Fig. 4d). Additionally, in order the further probe the proteomic alterations of levetiracetam in *App*^*NL-G-F/NL-G-F*^ mice, we performed a Bayesian analysis of variance in order to directly compare vehicle and levetiracetam treated *App*^*NL-G-F/NL-G-F*^ cohorts (Fig. S2a). Proteins that were identified as significantly upregulated with levetiracetam treatment were subjected to GO:CC enrichment analysis (Fig. S2b). This showed that GO:CC terms related to presynaptic endocytosis (e.g. HOPS complex, AP-2 adaptor complex, presynaptic endocytic zone) were once again revealed to have upregulated levels with levetiracetam treatment. To further investigate the possibility that the levetiracetam modulated proteins physically interact, we subjected the group of normalized proteins to STRING analysis and uncovered a robust protein-protein interaction hub (Fig. 4e). These regulated endocytic factors participate in all three predominant steps (i.e. initiation, assembly, and fission) suggesting the entire process of endocytosis is modulated by levetiracetam (Fig. 4f).

**Figure 4.**
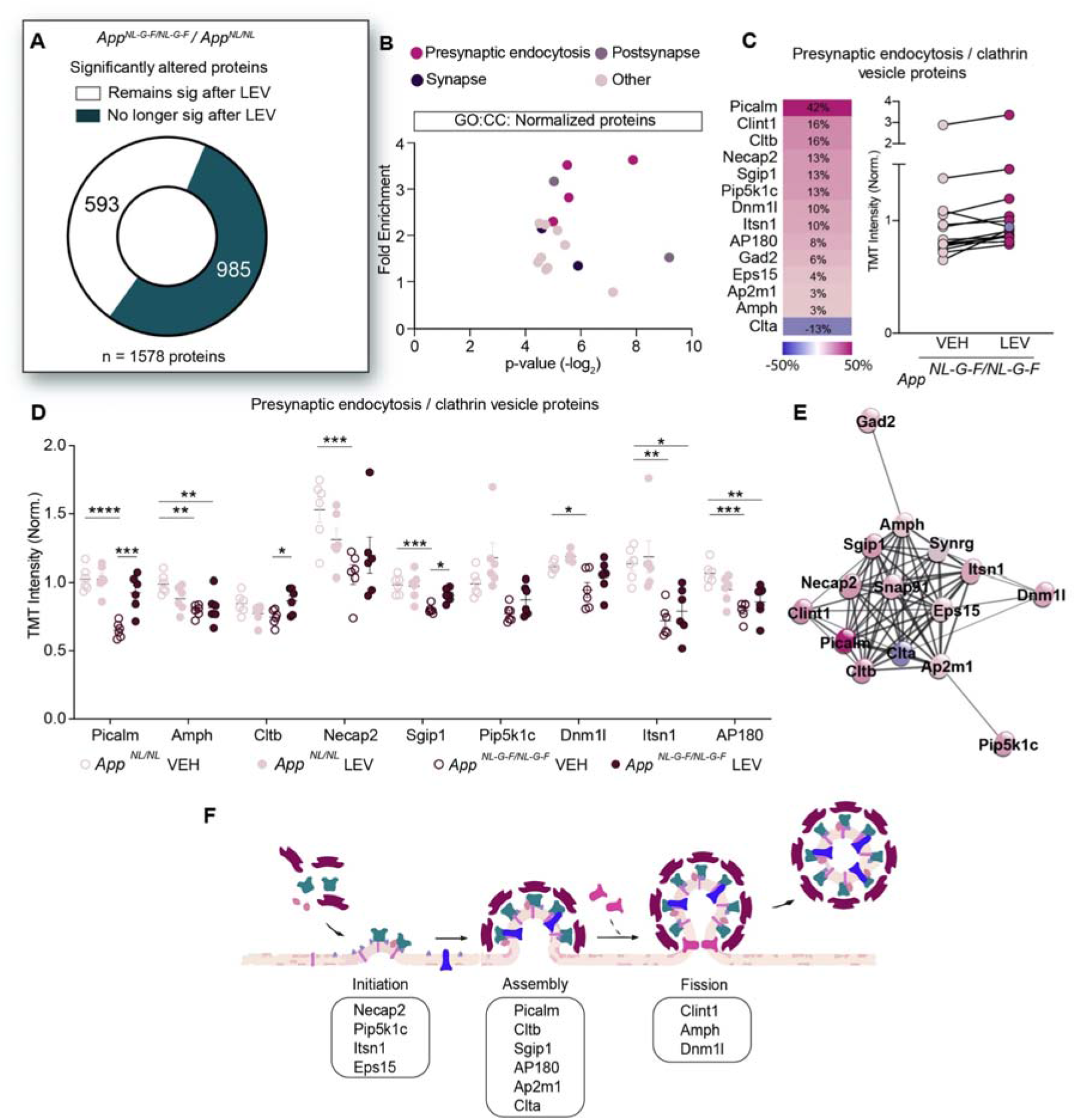
Chronic levetiracetam administration normalizes levels of presynaptic endocytic proteins in *App*^*NL-G-F/NL-G-F*^ cortex. (**A**) Pie chart depicts significantly altered proteins identified by comparing vehicle groups *App*^*NL-G-F/NL-G-F*^/*App*^*NL/NL*^. Dark green indicates the number of proteins that remain significantly altered after LEV. Light green denotes the proteins that no longer significantly altered after LEV (Table S3). (**B**) Gene ontology (GO) cell component enrichment (GO:CC) analysis plots depict fold enrichment versus p-value (−log_2_), analyzed by Fisher’s exact test. GO terms related to presynaptic endocytosis (pink), postsynapse (light purple), synapse (dark purple), and all other terms (light pink) (Table S4). (**C**) Percent change of presynaptic endocytosis proteins (GO:0098833) between VEH and LEV *App*^*NL-G-F/NL-G-F*^ groups. (**D**) Normalized presynaptic endocytosis protein abundance between *App*^*NL-G-F/NL-G-F*^ and *App*^*NL/NL*^, VEH and LEV groups. (**E**) Protein-protein interaction hub of presynaptic endocytosis proteins based on STRING functional enrichment analysis. **(F)** Cartoon depicting the stages of SV endocytosis. Data represents mean ± SEM, analyzed with unpaired Student’s t-test and BH correction. N=6 per genotype, N=6 per treatment group. Each circle represents an individual biological replicate. Data represents mean ± SEM, analyzed with unpaired Student’s t-test or one-way ANOVA with post-hoc Sidak test. * = p-value <.005.* = p-value <0.05, ** = p-value <0.01, *** = p-value < 0.001.

### Levetiracetam restores non-amyloidogenic APP processing in *App*^*NL-G-F/NL-G-F*^ and decreases Aβ_42_ levels

To further investigate the effect of levetiracetam treatment on amyloid deposition, we performed Thioflavin S staining on *App* KI sagittal sections. Thioflavin S puncta quantification revealed that treatment significantly decreased the amyloid plaque load in *App*^*NL-G-F/NL-G-F*^ cortex compared to vehicle treatment (p-value = 0.0046) (Fig. 5a-b). In line with previous literature, vehicle treated *App*^*NL-G-F/NL-G-F*^ had significantly more Aβ_42_ compared to vehicle *App*^*NL/NL*^ and *App*^*NL-F/NL-F*^ based on sandwich ELISA (p-value = <0.0001; p-value = <0.0001) (Fig. 5c). Interestingly, Aβ_42_ ELISA analysis revealed that levetiracetam treated *App*^*NL-G-F/NL-G-F*^ cortical extracts, have significantly reduced Aβ_42_ levels (p-value = 0.0010) compared to vehicle treated *App*^*NL-G-F/NL-G-F*^ (Fig. 5c). Since Aβ_42_ levels are reduced without altering levels of full-length APP protein, we investigated if levels of APP cleavage products, β-CTF and α-CTF, were altered by levetiracetam. Western blot analysis of β-CTF and α-CTF bands from cortical homogenates were quantified in both *App*^*NL/NL*^ and *App*^*NL-G-F/NL-G-F*^ vehicle and levetiracetam groups (Fig. 5d-g). Notably, analysis of the β-CTF / α-CTFs ratio from *App*^*NL-G-F/NL-G-F*^ mice indicated that levetiracetam significantly decreases β-CTFs and correspondingly increases α-CTFs compared to vehicle controls (p-value = 0.0010) (Fig. 5g). There was no significant difference between β-CTF / α-CTFs ratio in the two *App*^*NL/NL*^ groups. Importantly, in both *App*^*NL/NL*^ and *App*^*NL-G-F/NL-G-F*^, full length APP showed no change in abundance (p-value = 0.1117; p-value = 0.5334) (Fig. 5f). This finding suggests that chronic levetiracetam administration shifts APP processing towards the non-amyloidogenic pathway, which in turn limits Aβ_42_ production.

**Figure 5.**
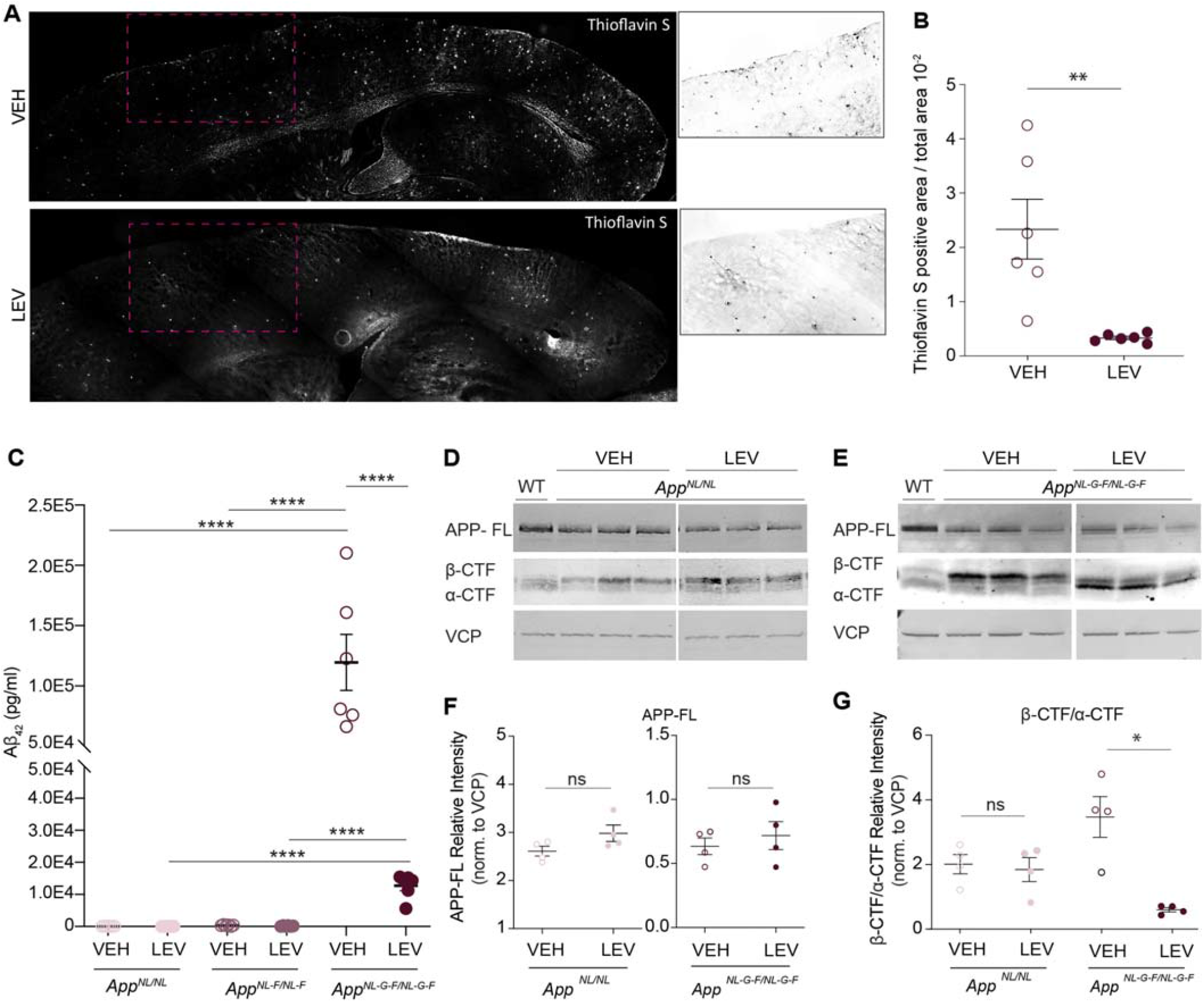
Chronic levetiracetam administration alters APP CTF production and decreases Aβ_42_ and in *App*^*NL-G-F/NL-G-F*^. (**A**) Cortical amyloid pathology in *App*^*NL-G-F/NL-G-F*^ comparing VEH and LEV treatment groups. Representative Thioflavin S stained sagittal brain sections are shown. White box indicates area of magnified image. (**B**) Quantification of amyloid plaque puncta normalized to cortical area. (**C**) Aβ_42_ levels in cortical homogenates from *App*^*NL/NL*^, *App*^*NL-F/NL-F*^, and *App*^*NL-G-F/NL-G-F*^ mice in VEH and LEV treated groups as measured by Aβ_42_ sandwich ELISA. N=6 for each genotype and treatment group. Each circle represents an individual biological replicate. (**D**) Representative WB analysis of full-length APP and APP cleavage products, β-CTF and α-CTF from cortical homogenates from *App*^*NL/NL*^ VEH and LEV groups. Age matched wild-type mouse age matched cortical homogenates were used as a negative control. VCP was used to control for loading and normalization. (**E**) Representative WB analysis of full-length APP and APP cleavage products, β-CTF and α-CTF from cortical homogenates from *App*^*NL-G-F/NL-G-F*^ groups. Age matched wild-type mouse age matched cortical homogenates were used as a negative control. VCP was used to control for loading. (**F**) Quantification of (D) showing the abundance of APP-FL normalized to VCP for *App*^*NL/NL*^ and *App*^*NL-G-F/NL-G-F*^, VEH and LEV groups. (**G**) Quantification of (E) showing the abundance of β-CTF/ α-CTF ratio normalized to VCP. N=4 for each genotype and treatment group. Each circle represents an individual biological replicate. Data represents mean ± SEM, analyzed with unpaired Student’s t-test or one-way ANOVA with post-hoc Sidak test. * = p-value <.05, ** = p-value < .01, *** = p-value < .001.

## DISCUSSION

Taken all together, our work shows that chronic levetiracetam treatment in *App* KI mouse models normalizes levels of presynaptic endocytosis machinery and alters APP proteolytic processing corresponding with lower levels of Aβ_42_ and decreased amyloid plaque deposits. As a growing body of evidence has demonstrated an association between AD and brain hyperexcitability, understanding the relationship between neural network dysfunction and Aβ pathology is crucial (Kuchibhotla et al., 2008, Bakker et al., 2012, Toniolo et al., 2020, Harris, et al., 2020, Sen et al. 2018). Interestingly, in a study of AD patients with epilepsy, a comparison of levetiracetam versus typical epilepsy drugs, lamotrigine and phenobarbital, demonstrated that while all the drugs were equally effective in reducing seizures, only levetiracetam treatment led to improved performance on cognitive tasks (Cumbo et al., 2010). Furthermore, in AD mouse models, only levetiracetam reduced hyperexcitability while also decreasing Aβ plaque burden and cognitive deficits (Vossel et al., 2017, Sanchez et al., 2012, Shi et al., 2013, Koh et al., 2010). This suggests that while hyperactivity contributes to increased Aβ pathology, treating hyperactivity alone is not sufficient to alleviate AD pathology. Our lab recently identified an impairment in turnover of SV associated proteins at early stages of AD pathology. In this study we hypothesized that levetiracetam’s unique beneficial effect on AD pathology could result from the atypical nature of this antiepileptic targeting the presynaptic SV2A protein (Hark et al., 2020). Our findings demonstrate that chronic levetiracetam treatment selectively normalizes levels of presynaptic endocytosis proteins and is capable of lowering Aβ_42_ levels by altering APP processing. Several supporting lines of evidence implicate dysregulation of endocytosis and presynaptic endocytic proteins in AD thus supporting why normalization of this process reduces amyloidogenic APP processing and ultimately Aβ_42_ production. Much of the previous evidence gathered on Aβ toxicity implicates the postsynaptic membrane as the primary site (DeBoer et al., 2014; Perdigão et al., 2020; Serrano-Pozo et al., 2011; Sheng et al., 2012). However, the localization and processing of APP mainly occurs at presynaptic terminals, and it has been previously shown that APP interacts with SVs (Masliah et al., 1994a; Oddo et al., 2003). Additionally, GWAS studies over the last decade have identified several AD-associated variants of endocytosis related genes including PICALM, BIN1, and SORL1 (Harold et al., 2009, Seshadri et al., 2014, Talwar et al., 2014). PICALM, which is a recruiter of adaptor complex 2 (AP-2) and is required for clathrin-mediated endocytosis, was the most significantly modulated protein in our datasets. How modulation of PICALM affects APP processing is not well understood. Some evidence supports an inverse relationship between PICALM levels and Aβ_42_ pathology. For example, APP^sw/0^ x PICALM^+/−^ mice displayed hippocampal and cortical Aβ loads 4-fold higher compared to APP^sw/0^ x PICALM^+/+^ controls (Zhao et al., 2015). In addition, it has been shown that AP-2 is required for APP endocytosis and has the ability to alter APP processing by promoting BACE1 trafficking (Bera et al., 2020). These studies proposed that AP-2 functions at the presynapse to sort BACE1 leading to a regulation of its degradation during neuronal activity (Bera et al., 2020). This would explain why rescuing levels of endocytosis proteins, such as AP-2, could result in a shift toward the non-amyloidogenic pathway of APP cleavage. Furthermore, in post-mortem AD brains, additional proteins functioning in endocytosis have reduced levels were also normalized in our datasets (e.g. AP180 and Dynamin1) (Yao et al., 1998, Yao et al., 2003). Taken all together, there is substantial evidence that suggests endocytosis and intracellular sorting determines how APP is processed. Our data supports the concept that levetiracetam lowers Aβ_42_ levels by normalizing the abundance of presynaptic endocytosis machinery that corresponds to a shift in APP processing towards the non-amyloidogenic pathway.

## ABBREVIATIONS

Aβ: amyloid-beta
APP: Amyloid Precursor Protein
AD: Alzheimer’s disease
*App* KI: APP knock-in
β-CTF/ α-CTF: beta/alpha carboxyl-terminal fragment
GO: Gene Ontology
LEV: levetiracetam
MS: mass spectrometry
SV: synaptic vesicle
SV2A: synaptic vesicle glycoprotein 2A
TMT: tandem mass tag
VEH: vehicle

## DECLARATIONS

### Ethical Approval and Consent to participate

Not applicable.

### Consent for publication

Not applicable.

### Availability of data and materials

Further information and requests for resources and reagents should be directed to and will be fulfilled by the Lead Contact, Jeffrey N Savas. The raw MS data will be deposited in MassIVE online database and upon acceptance will be available in the ProteomeXchange online database (http://www.proteomexchange.org/).

## Author Contributions

N.R.R. and J.N.S. designed the experiments; N.R.R. performed all the experiments; and N.R.R. and J.N.S. wrote the manuscript. All authors read and approved the final manuscript.

## Acknowledgments

We thank Tim Hark for contributing on Thioflavin S imaging and tissue collection. We thank the NU Center for Advanced Microscopy, which is generously supported by NCI CCSG P30 CA060553 awarded to the Robert H. Lurie Comprehensive Cancer Center.

## Authors’ information

Nalini R. Rao^1^ and Jeffrey N. Savas ^1^

^1^Department of Neurology, Northwestern University Feinberg School of Medicine, 303 E Chicago Ave, Ward 12-013, Chicago, IL 60611, USA.

## METHODS

### Animals

All experiments performed were approved by the Institutional Animal Care and Use Committee of Northwestern University (Protocols IS0009900 and IS00010858). The mice used for all experiments were Amyloid precursor protein Knock-in mice. These mice were originally obtained from the RIKEN Brain Science Institute, Saitama, Japan, from Dr. Takaomi C. Saido [Saito et al., 2014]. Mice were genotyped by Transnetyx using real-time PCR. For euthanasia, mice were anesthetized with isoflurane followed by cervical dislocation and acute decapitation. Equal numbers male and female mice were used for all experiments.

### Levetiracetam Injections and Brain Collection

Levetiracetam (United States Pharmacopeial) was dissolved in sterile saline solution (0.9% sodium chloride). Equal number male and female mice were randomly assigned to vehicle or treatment groups and were given chronic intra peritoneal injections of saline solution or 75mg/kg between 10am–1pm each day for 30 consecutive days. At the end of the 30 day chronic treatment, mice were anesthetized and transcardially perfused with cold PBS. Brains were then hemisected with one half for immunostaining and the other for biochemistry.

### TMT-MS Sample Preparation

TMT-MS sample preparation was performed as previously described [Jongkamonwiwat et al., 2020]. In brief, homogenized cortical brain extracts were prepared and 200µg of protein was used for TMT-MS sample preparation. Methanol-chloroform precipitation was used to separate proteins from lipids and impurities. Extracted protein was then resuspended in 6M guanidine in (100 mM HEPES). Proteins were further processed via the reduction of disulfide bonds with DTT, and alkylation of cysteine residues with iodoacetamide. Proteins were then digested for 3h at RT with 1μg LysC (Promega) and then digested overnight at 37°C with 2μg of Trypsin. The digest was then acidified with formic acid and desalted using C18 HyperSep columns (ThermoFisher Scientific). The eluted peptide solution was dried before resuspension in 100mM HEPES. Micro BCA assay was subsequently performed to determine the concentration of peptides. 100μg of peptide from each sample was then used for isobaric labeling. TMT 16-plex labeling was performed on peptide samples according to the manufacturer’s instructions (ThermoFisher Scientific). After incubating for 75 min at room temperature, the reaction was quenched with 0.3% (v/v) hydroxylamine. Isobaric labeled samples were then combined 1:1:1:1:1:1:1:1:1:1:1:1:1:1:1:1 and subsequently desalted with C18 HyperSep columns. The combined isobaric labeled peptide samples were fractionated into 8 fractions using High pH Reversed-Phase columns (Pierce). Peptide solutions were dried, stored at −80°C, and reconstituted in LC-MS Buffer A (5% acetonitrile, 0.125% formic acid) for LC-MS/MS analysis.

### TMT-MS Analysis

TMT-MS analysis was performed as previously described [Jongkamonwiwat et al., 2020]. In short, samples were resuspended in 20µl Buffer A (5% acetonitrile, 0.125% formic acid) and micro BCA was performed. 3µg of each fraction was loaded for LC-MS analysis via an auto-sampler with a Thermo EASY nLC 100 UPLC pump onto a vented Pepmap100, 75um x 2 cm, nanoViper trap column coupled to a nanoViper analytical column (Thermo Scientific) with stainless steel emitter tip assembled on the Nanospray Flex Ion Source with a spray voltage of 2000V. Orbitrap Fusion was used to generate MS data. The chromatographic run was performed with a 4h gradient beginning with 100% Buffer A and 0% B and increased to 7% B over 5 mins, then to 25% B over 160 mins, 36% B over 40 mins, 45% B over 10 mins, 95% B over 10 mins, and held at 95% B for 15 mins before terminating the scan. Buffer A contained 5% ACN and 0.125% formic acid in H20, and Buffer B contained 99.875 ACN with 0.125% formic acid. Multinotch MS3 method was programmed as the following parameter: Ion transfer tube temp = 300 °C, Easy-IC internal mass calibration, default charge state = 2 and cycle time = 3 s. MS1 detector set to orbitrap with 60 K resolution, wide quad isolation, mass range = normal, scan range = 300–1800 m/z, max injection time = 50 ms, AGC target = 6 × 10^5^, microscans = 1, RF lens = 60%, without source fragmentation, and datatype = positive and centroid [Jongkamonwiwat et al., 2020]. Monoisotopic precursor selection was set to included charge states 2–7 and reject unassigned. Dynamic exclusion was allowed n = 1 exclusion for 60 s with 10ppm tolerance for high and low. An intensity threshold was set to 5 × 103. Precursor selection decision = most intense, top speed, 3s. MS2 settings include isolation window = 0.7, scan range = auto normal, collision energy = 35% CID, scan rate = turbo, max injection time = 50 ms, AGC target = 6 * 10^5^, Q = 0.25. In MS3, the top ten precursor peptides were selected for analysis were then fragmented using 65% HCD before orbitrap detection. A precursor selection range of 400–1200 m/z was chosen with mass range tolerance. An exclusion mass width was set to 18 ppm on the low and 5 ppm on the high. Isobaric tag loss exclusion was set to TMT reagent. Additional MS3 settings include an isolation window = 2, orbitrap resolution = 60 K, scan range = 120 – 500 m/z, AGC target = 6*10^5^, max injection time = 120 ms, microscans = 1, and datatype = profile.

### TMT-MS Data Analysis and Quantification

TMT-MS data analysis was performed as previously described in [Jongkamonwiwat et al., 2020]. In short, protein identification, TMT quantification, and analysis were performed with The Integrated Proteomics Pipeline-IP2 (Integrated Proteomics Applications, Inc., http://www.integratedproteomics.com/). Proteomic results were analyzed with ProLuCID, DTASelect2, Census, and QuantCompare. MS1, MS2, and MS3 spectrum raw files were extracted using RawExtract 1.9.9 software (http://fields.scripps.edu/downloads.php). Pooled spectral files from all eight fractions for each sample were then searched against the Uniprot mouse protein database and matched to sequences using the ProLuCID/SEQUEST algorithm (ProLuCID ver. 3.1) with 50 ppm peptide mass tolerance for precursor ions and 600 ppm for fragment ions. Fully and half-tryptic peptide candidates were included in search space, all that fell within the mass tolerance window with no miscleavage constraint, assembled and filtered with DTASelect2 (ver. 2.1.3) through the Integrated Proteomics Pipeline (IP2 v.5.0.1, Integrated Proteomics Applications, Inc., CA, USA). Static modifications at 57.02146 C and 304.2071 K and N-term were included. The target-decoy strategy was used to verify peptide probabilities and false discovery ratios [McAlister et al., 2014]. Minimum peptide length of five was set for the process of each protein identification and each dataset included a 1% FDR rate at the protein level based on the target-decoy strategy. Isobaric labeling analysis was established with Census 2 as previously described TMT channels were normalized by dividing it over the sum of all channels [McAlister et al., 2014]. No intensity threshold was applied. The fold change was then calculated as the mean of the experimental group standardized values and p-values were then calculated by Student’s t-test with Benjamini-Hochberg adjustment.

### Online Databases for PANTHER and STRING (http://string-db.org)

Protein ontologies were determined with protein analysis through evolutionary relationships (PANTHER) system (http://www.pantherdb.org), in complete cellular component categories [Mi et al., 2016]. The statistical overrepresentation test was calculated by using the significant proteins identified from comparing VEH versus LEV experimental groups for each *App* KI genotype as the query and the aggregated total proteins identified in all three comparisons as the reference. Protein ontologies with Fisher statistical tests with false discovery rate (FRD) correction less than 0.05 were considered significant.

The Search Tool for the Retrieval of Interacting Genes (STRING) database was used to determine protein-protein interactions from significant quantified proteins identified by GO:CC term. The STRING resource is available at http://string-db.org [Szklarczyk et al., 2017]. The corresponding protein–protein interaction networks were constructed with a highest confidence of interaction score at 0.9.

### Thioflavin Staining

After transcardial perfusion with cold PBS, hemisected brains were drop fixed in 4% PFA overnight, cryoprotected in 30% sucrose for two days, embedded in a cryomold with OCT, flash frozen on dry ice and stored at −80°C until cryosectioning. 30µm sagittal cryosections were prepared and mounted onto gelatin coated slides (SouthernBiotech). Section were then prepared for thioflavin S staining following standard procedures [Ly et al., 2011]. In short, sections were washed with 70% ethanol for one minute followed by 80% ethanol for one minute before being incubated in filtered thioflavin S solution (1% in 80% ethanol) for 15 minutes in the dark. Slides were then washed sequentially with 80% then 70% ethanol then distilled water for one minute each. Coverslips were mounted using Fluoromount-G (SouthernBiotech). Sections were imaged at the Northwestern University Center for Advanced Microscopy with a TissueGnostics system using a 10X objective. Analysis was conducted using Fiji with the Analyze puncta tool following thresholding. Cortical area analyzed was kept consistent throughout each section.

### Aβ_42_ ELISA Assay

Aβ_42_ levels were measured using a human Aβ_42_ ELISA kit (Thermo Scientific) following manufacturer instructions. In short, 5M guanidine HCl was added to cortical homogenates (1-2 mg) and kept shaking for 1 hour at RT. Samples were then diluted 1:10 for *App*^*NL/NL*^ and *App*^*NL-F/NL-F*^ and 1:1000 for *App*^*NL-G-F/NL-G-F*^ in Standard Diluent buffer. 50μl of sample was loaded into wells coated with provided Aβ_42_ antibody and incubated for 3 hours at RT. After 3 washes, HRP conjugated antibody was added for 30 minutes. After another wash step, samples were incubated with stabilized chromogen for 30 minutes, and the reaction was stopped with an acid-based Stop solution. Finally, OD was measured at 450 nm using a Synergy HTX multi-mode microplate reader (Biotek) and compared to a standard curve to determine the final concentration.

### Western Blotting

Cortical brain extracts were homogenized in 500µl homogenization buffer (4 mM HEPES, 0.32 M sucrose, 0.1 mM MgCl2) supplemented with a protease inhibitor cocktail (aprotinin, leupeptin, AEBSF, benzamidine, PMSF, and pepstatin A. Tissue was then homogenized using a bead based Precellys homogenizer. Protein concentration was then determined by BCA assay (Thermo Scientific) per manufacturer’s instructions and compared with the respective standard curve. 50µg of each sample was then prepared for western blots by adding 6X SDS sample buffer. The mixtures were sonicated and boiled at 96°C for 5 minutes each and then loaded in a 16% Tris-glycine gel. Gels were run at 80 V for 4 hours, then were wet transferred to a 0.2μm nitrocellulose membrane. Membranes were then blocked with Odyssey Blocking Buffer (LI-COR) in PBS for 1 hour then incubated overnight with Anti-Amyloid beta precursor protein (Y188) rabbit monoclonal antibody at 1:1,000 (Abcam Cat# ab32136) and Anti-VCP mouse monoclonal antibody at 1:2,000 (Abcam Cat# ab11433). The next day membranes were washed and incubated in secondary antibody IRDye 800CW Donkey anti-Rabbit IgG antibody (LI-COR Biosciences Cat# 926-32213) and IRDye 680RD Donkey anti-Mouse IgG antibody (LI-COR Biosciences Cat# 925-68072) for 1 hour at RT. Blots were imaged on an Odyssey CLx (LI-COR).

### Quantification and Statistical Analysis

Statistical analyses were performed using GraphPad Prism. All values in figures with error bars are presented as mean ± SEM. Comparisons of vehicle versus levetiracetam groups was performed using unpaired Student’s t-tests. Comparisons across all three genotypes were compared by one-way ANOVA and post-hoc Fisher’s test. P-values < 0.05 were considered statistically significant. Multiple test correction was performed with the Benjamini-Hochberg correction. For Bayesian analysis of variance, we implemented BAMarray 2.0, a Java software package that implements the Bayesian ANOVA for microarray (BAM) algorithm (Ishwaran et al., 2003). The BAM approach uses a special type of inferential regularization known as spike-and-slab shrinkage, which provides an optimal balance between total false detections (the total number of genes falsely identified as being differentially expressed) and total false non-detections (the total number of genes falsely identified as being non-differentially expressed) (Ishwaran et al., 2003).

## FIGURE LEGNEDS

**Figure S1.**
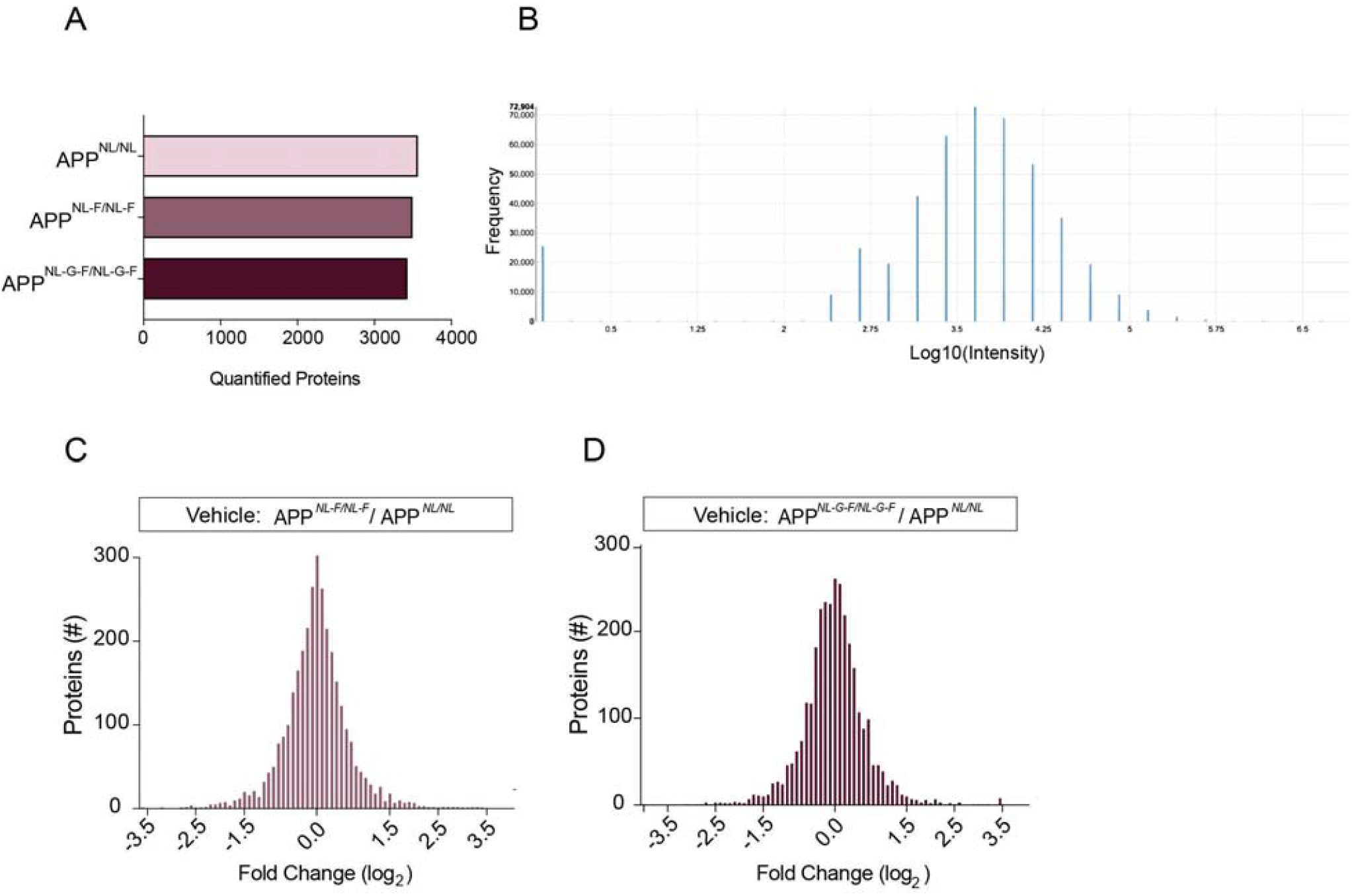
TMT-MS experiments have similar data quality. (**A**) Total quantified proteins for each *App*^*NL/NL*^, *App*^*NL-F/NL-F*^, and *App*^*NL-G-F/NL-G-F*^ TMT-MS analyses. (**B**) Representative reporter ion intensity distribution of 16-plex TMT labelling. (**C**) Global cortical proteome remodeling of vehicle *App*^*NL-F/NL-F*^ /*App*^*NL/NL*^. (**D**) Global cortical proteome remodeling of vehicle *App*^*NL-G-F/NL-G-F*^ /*App*^*NL/NL*^. N=6 mice for each group.

**Figure S2.**
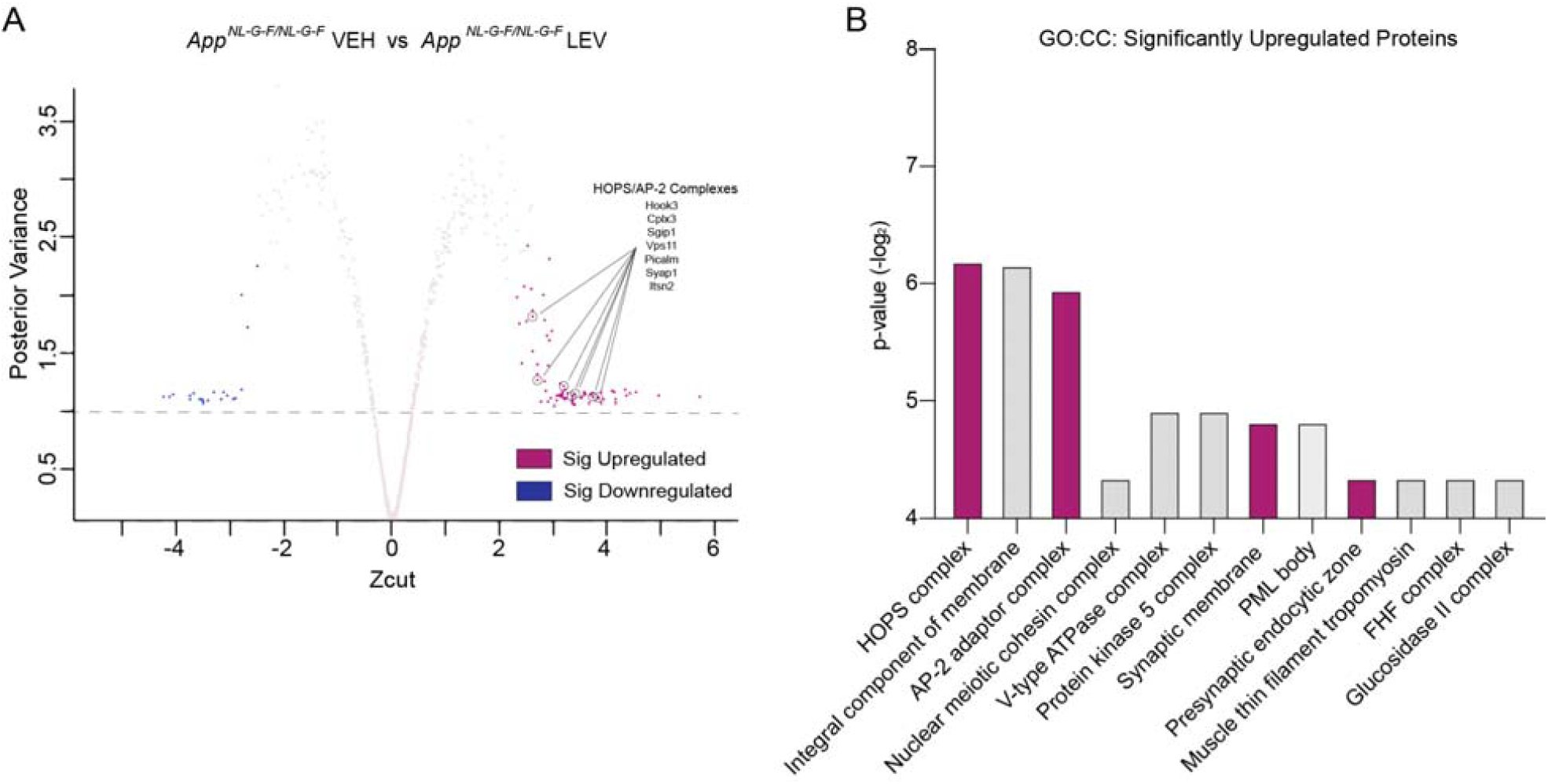
Presynaptic endocytic proteins are significantly upregulated with levetiracetam treatment in *App*^*NL-G-F/NL-G-F*^. (**A**) Proteins that are differentially expressed using Bayesian ANOVA when directly comparing vehicle and levetiracetam treatment in *App*^*NL-G-F/NL-G-F*^ cohorts. Pink and blue dots indicated significantly up and down regulated proteins respectively. Grey dots indicate non-significant proteins. (**B**) Gene ontology (GO) cell component enrichment (GO:CC) analysis of significantly upregulated proteins plots depict p-value (−log_2_) for each GO:CC term.

Table S1

Summary of relative abundances for significantly differentially regulated proteins across *APP* KI genotypes from cortical extracts. Related to Figure 2. Each tab represents a dataset. Experiment column indicates the genotype comparison. The average relative TMT intensities are shown for all significant proteins. T-test p-values and BH adjusted p-values are indicated.

Table S2

Significantly overrepresented cellular components from a GO analysis using the query of proteins that were significantly differentially regulated in each dataset against the reference of all proteins identified. Related to Figure 2. Each tab represents a dataset. Fold enrichment and p-values were obtained from the Fisher’s exact test are shown.

Table S3

Summary of significantly altered proteins identified by comparing vehicle groups *App*^*NL-G-F/NL-G-F*^ */App*^*NL/NL*^ and the proteins that either remain or no longer are significantly altered after LEV. Related to Figure 4. The average relative TMT intensities are shown for each group. T-test p-value and BH adjusted p-values are indicated.

Table S4

Significantly overrepresented cellular components from a GO analysis using the query of proteins that were no longer significantly differentially regulated after LEV against the reference of all proteins identified. Related to Figure 4. Each tab represents a dataset. Fold enrichment and p-values were obtained from the Fisher’s exact test are shown.

